# Physiological and transcriptional profiling of surfactin exerted antifungal effect against *Candida albicans*

**DOI:** 10.1101/2022.04.19.488861

**Authors:** Ágnes Jakab, Fruzsina Kovács, Noémi Balla, Zoltán Tóth, Ágota Ragyák, Zsófi Sajtos, Kinga Csillag, Csaba Nagy-Köteles, Dániel Nemes, István Pócsi, László Majoros, Ákos T. Kovács, Renátó Kovács

**Affiliations:** Department of Molecular Biotechnology and Microbiology, Institute of Biotechnology, Faculty of Science and Technology, University of Debrecen, Debrecen, Hungary; Department of Medical Microbiology, Faculty of Medicine, University of Debrecen, Debrecen, Hungary; Doctoral School of Pharmaceutical Sciences, University of Debrecen, Debrecen, Hungary; Department of Inorganic and Analytical Chemistry, Agilent Atomic Spectroscopy Partner Laboratory, University of Debrecen, Debrecen, Hungary; Department of Pharmaceutical Technology, Faculty of Pharmacy, University of Debrecen, Debrecen, Hungary; Bacterial Interactions and Evolution Group, DTU Bioengineering, Technical University of Denmark, Kongens Lyngby, Denmark; Faculty of Pharmacy, University of Debrecen, Debrecen, Hungary

**Keywords:** *Candida albicans*, surfactin, probiotic, transcriptomics, metal, *Bacillus subtilis*

## Abstract

Given the risk of *Candida albicans* overgrowth in the gut, novel complementary therapies should be developed to reduce fungal dominancy. This study highlights the antifungal characteristics of a *Bacillus subtilis*-derived secondary metabolite, surfactin with high potential against *C. albicans*. Surfactin inhibited the growth of *C. albicans* following a 1-hour exposure, in addition to reduced adhesion and morphogenesis. Specifically, surfactin did not affect the level of reactive oxygen species but increased the level of reduced glutathione. Surprisingly, ethanol production enhanced following 2 hours of surfactin exposure. Surfactin treatment caused a significant reduction in intracellular iron, manganese and zinc content compared to control cells, whereas the level of copper was not affected. Alongside these physiological properties, surfactin also enhanced fluconazole efficacy. To gain detailed insights into the surfactin-related effects on *C. albicans*, genome-wide gene transcription analysis was performed. Surfactin treatment resulted in 1390 differentially expressed genes according to total transcriptome sequencing (RNA-Seq). Of these, 773 and 617 genes with at least a 1.5-fold increase or decrease in transcription, respectively, were selected for detailed investigation. Several genes involved in morphogenesis or related to metabolism (e.g., glycolysis, fermentation, fatty acid biosynthesis) were down-regulated. Moreover, surfactin decreased the expression of *ERG1*, *ERG3*, *ERG9*, *ERG10* and *ERG11* involved in ergosterol synthesis, whereas genes associated with ribosome biogenesis and iron metabolism and drug transport-related genes were up-regulated. Our data demonstrate that surfactin significantly influences the physiology and gene transcription of *C. albicans*, and could contribute to the development of a novel innovative complementary therapy.

**Importance:** Although gut colonization by *Candida albicans* can be considered normal, it may be associated with intestinal diseases. Furthermore, *Candida* dominance in the gut may pose a potent risk for systemic candidiasis, especially for immunocompromised individuals. In recent years, interest has been growing for the use of *Bacillus subtilis* as a safe and effective probiotic for human healthcare. Surfactin is a *B. subtilis*-derived lipopeptide with potential antifungal activity; however, the mechanism underlying this remains unknown. In this study, surfactin negatively affected the adherence, morphogenesis and metabolism of *C. albicans* and increased ethanol production. These were associated with a reduction in intracellular iron, manganese and zinc while the copper content was not affected. Alongside these physiological modulations, surfactin also had a potent synergistic effect on fluconazole. Our results provide a definitive explanation for the surfactin-related antifungal effect of *B. subtilis*; furthermore, these data provide a good basis for future probiotic development.

## Introduction

Molecular typing of *Candida albicans* isolates showed high similarity between those isolated from blood and isolates colonizing the gastrointestinal tract proving that gut might be one of the most important sources of invasive candidiasis (1–2). Microbial dysbiosis and *Candida* dominance in the gut can be considered as a potent risk for systemic candidiasis, especially in patients treated with cytotoxic agents that disrupt the intestinal mucosa (3). Thus, it is important to develop and introduce new supportive and complementary therapeutic strategies in order to reduce the overgrowth of *Candida* cells in the gut. The administration of various probiotics may be a useful approach in this context (4–6). Different species from the *Lactobacillus* and *Bifidobacterium* genera are the most frequently used probiotics. Nevertheless, other potential probiotic species have also been shown to be effective for the treatment of gastrointestinal candidiasis (7).

In the last decade, there has been increasing interest in the use of *Bacillus subtilis* strains as safe and effective probiotics for human healthcare (7). On the one hand, *B. subtilis* spores modulate the immune response of the host; on the other hand, the vegetative form can release enzymes, antioxidants and vitamins to support digestion. It is important to emphasize that *B. subtilis*-derived exopolysaccharides have relevant immunomodulatory effect, which may prevent inflammatory disease induced by enteric pathogens (8–9). Moreover, several strains of *B. subtilis* can secrete antimicrobial compounds, promoting the optimal microbial balance (10–12). Generally, members of the *B. subtilis*-species complex are potent source for natural products with anti-fungal compounds, including anti-*Candida* activity (13). Among these natural products, surfactin is a *B. subtilis*-derived cyclic lipopeptide that shows activity against gram-positive bacteria and has a wide range of targets, so it is difficult for microorganisms to develop drug resistance against it (10–12). Previous studies showed that its oral half-lethal dose in mice exceeded 5000 mg/kg, and that acute toxicity was low or lacking (14).

Recent studies revealed that surfactin has an effect on yeasts; furthermore, it effectively decreases the adhesion of *C. albicans* and its hydrophobicity (15–18). Nevertheless, the effect of surfactin on *C. albicans* has not been sufficiently investigated and the underlying mechanisms of surfactin-induced effects remain unknown. Understanding how surfactin can influence the physiological and genetic properties of *C. albicans* would be pivotal in further probiotic development. To gain further insights into its previously observed inhibitory potential, we report the effect of surfactin on cell growth, adhesion, virulence, metabolism, and intracellular metal content, in addition to the genome-wide gene expression changes caused by surfactin exposure as determined using total transcriptome sequencing (RNA-Seq).

## Materials and methods

### Fungal strain, culture media and epithelial cell line

The SC5314 *C. albicans* wild-type reference strain was maintained and cultured on yeast extract-peptone-dextrose (YPD) (VWR, 2% glucose, 2% peptone, 1% yeast extract with or without 2% agar) as described previously (19). Susceptibility experiments to fluconazole (Merck, Budapest, Hungary) and surfactin was determined in RPMI-1640 (with L-glutamine and without bicarbonate, pH 7.0 with MOPS; Merck, Budapest, Hungary). For toxicity and adhesion experiments, the Caco-2 epithelial cell line was obtained from the European Collection of Cell Cultures (ECACC) (Salisbury, United Kingdom) and cultured as described previously by Nemes et al. (20). Briefly, cells were grown in plastic cell culture flasks in Dulbecco’s modified Eagle’s medium (DMEM) supplemented with 3.7 g/l NaHCO_3_, 10% (vol/vol) heat-inactivated fetal bovine serum, a 1% (vol/vol) non-essential amino acid solution, 0.584 g/l L-glutamine, 4.5 g/l D-glucose, 100 IU/ml penicillin, and 100 mg/l streptomycin at 37°C in the presence of 5% CO_2_. The cells were maintained by regular passaging, and glutamine was supplemented with GlutaMAX. Cells between passages 20 and 40 were used for adhesion and toxicity experiments (20).

### Toxicity experiments

Concentrations of 16 mg/l, 32 mg/l, 64 mg/l, 128 mg/l and 256 mg/l surfactin were evaluated regarding toxicity to Caco-2 cell line using a 3-(4,5-dimethyl-2-thiazolyl)-2,5-diphenyl-2H-tetrazolium bromide (MTT) assay (Merck, Budapest, Hungary) and xCELLigence real-time cell analysis (ACEA Biosciences, Inc., San Diego, CA, USA), where none of them caused relevant toxicity (19–20). A final concentration of 128 mg/l surfactin (Merck, Budapest, Hungary) was used in our experiments, corresponding to the average surfactin production of various *B. subtilis* strains (21).

### Growth assay

*C. albicans* pre-cultures were grown in 5 ml YPD medium at 37°C for 18 hours, diluted to an optical density of 0.1 (OD_640_) with YPD then grown further at 37°C and at 2.3 Hz shaking frequency. Following a 4-hour incubation period, some cultures were supplemented with 128 mg/l surfactin, and microbial growth was followed by measuring changes in optical density at 640 nm and dry cell mass (DCM) (19, 22). Samples were taken at 1 hour for RNA isolation and for measurement of redox changes, phospholipase, lipase and extracellular protease activity. Two- and 4-hour surfactin exposures were used to determine glucose consumption and ethanol production. The glutathione and intracellular metal contents were determined following 4 hours of surfactin treatment.

Morphological changes exerted by surfactin were examined after 4, 5, 6 and 8 hours of incubation at 37 °C using a Zeiss Axioskop 2 mot microscope coupled with a Zeiss Axiocam HRc camera using the phase-contrast technique (23).

For all experiments, statistical analyses were performed between control and surfactin-treated cultures using the paired Student’s *t* test. Significance was defined as a *p* value of < 0.05.

### Phospholipase, esterase and extracellular protease activity

Extracellular phospholipase production by surfactin-treated (128 mg/l) and untreated *C. albicans* cells was investigated on egg yolk medium [4% glucose, 1% peptone, 5.85% NaCl, 0.05% CaCl_2_, and 10% (vol/vol) sterile egg yolk emulsion, 1.5% agar]. Esterase activity was determined on Tween-20 medium [1% peptone, 0.5% NaCl, 0.01% CaCl_2_×2H_2_O, 1% (vol/vol) Tween-20 (Merck, Budapest, Hungary), 2% agar]. Aspartic proteinase secretion was evaluated on solid bovine serum albumin medium [0.02% MgSO_4_×7H_2_O, 0.25% K_2_HPO_4_, 0.5% NaCl, 0.1% yeast extract, 2% glucose and 0.25% bovine serum albumin (Merck, Budapest, Hungary), 2% agar].

To assess the production of virulence-related enzymes, 5-µl suspensions of *Candida* cells (1×10^7^ CFU/ml) were inoculated onto agar plates as described previously (24). Colony diameter and precipitation zones on egg yolk and Tween-20 plates and clear areas on bovine serum albumin media (Pz) were measured after 7 days of incubation at 37°C (22). Enzyme activities were measured in three independent experiments and are presented as means with standard deviations.

### Determination of glucose consumption and ethanol production

Aliquots of *C. albicans* culture media were collected by centrifugation (5 min, 4000×g, 4°C) following 2 and 4 hours of surfactin exposure. Changes in the glucose contents of the *C. albicans* supernatants were determined using the glucose oxidase assay described by Jakab et al. (25).

The concentration of ethanol in culture media was determined by headspace gas chromatography (HS-GC-FID system, PerkinElmer GC, Clarus 680 with PerkinElmer TurboMatrix 40 Trap Headspace Sampler, FID with helium as the carrier gas 1 ml/min) as described previously (25–26). Briefly, static HS injection was performed with a 1-µl injection volume, and a capillary column (Agilent, DB×5.625, 30 m×0.25 mm I.D.) was used for separation. Data analysis was performed with the PerkinElmer TotalChrom Workstation V.6.3.2 Data System.

Glucose consumption and ethanol fermentation values were calculated and expressed in DCM units (mg/kg). The glucose consumption and ethanol production by dry biomasses were determined in three independent experiments and mean±standard deviation values were calculated.

### Determination of intracellular metal contents of C*andida albicans* cells

The culturing of fungal cells and surfactin treatment was performed as described above. *Candida* cells were collected by centrifugation (5 min, 4 000×g, 4°C) after 4 hours of incubation following surfactin exposure. The intracellular metal (Fe, Zn, Cu, and Mn) contents of the dry biomass were measured by inductively coupled plasma optical emission spectrometry (ICP-OES; 5110 Agilent Technologies, Santa Clara, CA, USA) following atmospheric wet digestion in 3 ml of 65% HNO_3_ and 1 ml of 30% H_2_O_2_ in glass beakers. The metal contents of the samples were calculated in DCM units (in milligrams per kilogram) using the method reported by Jakab et al. (23). The metal contents of the dry biomass were determined in triplicate and mean±standard deviation values were calculated.

### Reactive species and glutathione production caused by surfactin exposure

Reactive species were determined with or without 1 hour of surfactin exposure (128 mg/l) by a reaction that converts 2′,7′-dichlorofluorescin diacetate to 2′,7′-dichlorofluorescein (DCF) (Sigma, Budapest, Hungary) (19). The amount of DCF produced is proportional to the quantity of reactive species.

Following a 4-hour surfactin exposure, reduced glutathione (GSH) and oxidized glutathione (GSSG) contents were measured with 5,5′-dithio-bis (2-nitrobenzoic acid)-glutathione reductase assay in cell-free yeast extracts prepared by 5-sulfosalicylic acid treatment as described by Emri et al. (27).

Reactive species, GSH and GSSG were measured in three independent experiments and are presented as means±standard deviation.

### Adherence of *Candida albicans* cells to inert surface and the metabolic activity of adhered fungal cells over time

The density of the *Candida* suspension used in this experiment was 1×10^6^ CFU/ml. A total of 100 μl of the *C. albicans* suspension was pipetted into 96-well flat-bottom polystyrene microtitre plates (TPP, Trasadingen, Switzerland) in the presence of 128 mg/l surfactin and incubated statically at 37°C. Pre-determined wells were assigned to endpoints of 2, 4, 6, 8, 10, 12 and 24 hours. After 2, 4, 6, 8, 10, 12 and 24 hours, the corresponding pre-assigned wells were washed three times with sterile physiological saline then the metabolic activity of adhered cells was measured using the XTT-assay as described previously (19). The percent change in metabolic activity was calculated on the basis of the absorbance (*A*) at 492 nm as 100%×(*A*_well_–*A*_background_)/(*A*_drug-free well_–*A*_background_). Three independent experiments were performed and the mean ± standard deviation was calculated for each examined time point. Statistical comparisons of relative metabolic activity data were performed using the paired Student’s *t* test. The differences between values for treated and control cells were considered significant if the *p* value was <0.05.

### Adherence of *Candida albicans* cells to Caco-2 epithelial cells

For adhesion-related experiments, 1×10^5^ epithelial cells were seeded on glass coverslips (pre-treated with rat tail derived collagen I; Gibco—Thermo Fisher, USA), placed in 24-well plates and incubated for 3 days prior to infection (28). For infection, pre-cultured *C. albicans* cells were collected by centrifugation (5 min, 4000×g, 4°C), washed three times with sterile physiological saline and adjusted to 2×10^5^ cells/ml in DMEM. Fungal cells were co-incubated with Caco-2 epithelial cells for 1 hour in serum-free DMEM with or without 128 mg/l surfactin at 37°C in the presence of 5% CO_2_. After co-incubation of epithelial cells with *C. albicans* cells, non-adherent yeast cells were removed by washing three times with sterile physiological saline and the cells on the cover slips were fixed with 4% formaldehyde. Adherent fungal cells were stained with calcofluor white (CFW; Merck, Budapest, Hungary) and were visualized with a fluorescence microscope (Zeiss Axioskop 2 mot microscope coupled with a Zeiss Axiocam HRc camera) (28). The number of adherent cells was determined by counting at least 50 fields. Adhesion (%) was calculated using the following formula: [(average cell count in the field area of the well in μm^2^)/(area of the field in μm^2^ inoculated yeast cells in each well)]×100 (28). Statistical analysis was performed between control and surfactin-treated cultures using the paired Student’s *t* test. Significance was defined as a *p* value of < 0.05.

### Interactions between fluconazole and surfactin *in vitro*

Interactions between fluconazole and surfactin were examined using a two-dimensional broth microdilution chequerboard assay. Interactions were analyzed by calculating the fractional inhibitory concentration index (FICI) and using the Bliss independence model (29). Interactions were determined in three independent experiments. The tested concentrations ranged from 0.004 to 1 mg/l for fluconazole and from 4 to 256 mg/l for surfactin. FICIs were determined using the following formula: ΣFIC=FIC_A_+FIC_B_=[(MIC_A_^comb^/MIC_A_)]+[(MIC_B_ /MIC_B_)], where MIC_A_ and MIC_B_ stand for the MICs of drugs A and B when used alone, and MIC_A_ and MIC_B_ represent the MICs of drugs A and B in combination at isoeffective combinations, respectively. FICIs were determined as the lowest ΣFIC. The MICs of the drugs alone and of all combinations were determined as the lowest concentration resulting in at least a 50% decrease in turbidity compared to the unexposed control cells. If the obtained MIC was higher than the highest tested drug concentration, the next highest twofold concentration was considered the MIC value. FICIs ≤ 0.5 were defined as synergistic, between > 0.5 and 4 as indifferent, and > 4 as antagonistic (19, 29).

Fluconazole and surfactin interactions were further analyzed using the Bliss independence model with help of the open access Synergyfinder ® application. Briefly, in this software, the Bliss independence model compares the observed versus predicted inhibition response, where a mean synergy score of less than −10 is considered antagonistic, between −10 and 10 is considered to be indifferent and above 10 is synergistic (30).

### RNA sequencing

For RNA extraction, fungal cells were collected 1 hour following surfactin exposure by centrifugation (5 minutes, 4,000×g at 4°C). Fungal cells were washed three times with phosphate-buffered saline (PBS) and stored at −70°C until examination. Total RNA samples were prepared from lyophilized cells (CHRIST Alpha 1-2 LDplus lyophilizer, Osterode, Germany) derived from untreated and surfactin-exposed cultures (22–23). Three independent cultures were used for RNA-seq experiments and RT-qPCR tests. The quality of RNA was determined using the Eukaryotic Total RNA Nano kit (Agilent, USA) in an Agilent Bioanalyzer. RNA-Seq libraries were prepared from total RNA using an Ultra II RNA Sample prep kit (New England BioLabs) according to the manufacturer’s protocol. The single read 75-bp sequencing reads were generated on an Illumina NextSeq500 instrument. Approximately 23–27 million reads were generated per sample. RNA-Seq libraries were made by the Genomic Medicine and Bioinformatics Core Facility of the Department of Biochemistry and Molecular Biology, Faculty of Medicine, University of Debrecen, Hungary. Raw reads were aligned to the reference genome (https://www.ncbi.nlm.nih.gov/genome/?term=Candida+albicans+SC5314), and the percentage of aligned reads varied between 92% and 95% in each sample. Gene transcription differences between surfactin-treated and control groups were compared using a moderated *t* test; the Benjamini-Hochberg false discovery rate was used for multiple-testing correction, and a corrected *P* value of <0.05 was considered significant. Up- and down-regulated genes were defined as differentially expressed genes with >1.5-fold change (FC, up-regulated genes) or less than −1.5-FC (down-regulated genes). The FC ratios were calculated from the normalized gene transcription values.

### Reverse transcription quantitative real-time PCR (RT-qPCR) assays

Surfactin-induced changes in the transcription of certain membrane transport, virulence and primary metabolism genes were validated by RT-qPCR (22–23). The RT-qPCRs were performed with a Luna Universal one-step RT-qPCR kit (NEB, USA) according to the manufacturer’s protocol using 500 ng of DNase (Merck, Budapest, Hungary)-treated total RNA per reaction. Oligonucleotide primers (Table S1) were designed using Oligo Explorer (version 1.1.) and Oligo Analyzer (version 1.0.2) software. Three parallel measurements were performed for every sample in a LightCycler 96 real-time PCR instrument (Roche, Switzerland). Relative transcription levels (ΔΔCP value) were determined based on the following formula: ΔCPcontrol−ΔCPtreated where ΔCPcontrol=CPtested gene, control−CPreference gene, control for untreated cultures, while ΔCP_treated_=CP_tested gene, treated_−CP_reference gene, treated_ for surfactin-treated cultures. The CP values represent the RT-qPCR cycle numbers of crossing points. Two reference genes (*TUB1* and *HPT1*) were tested. All showed stable transcription in our experiments and, hence, only data relative to the *HPT1* (C2_02740C) transcription values are presented.

### Gene set enrichment analysis

Gene set enrichment analyses on the up-regulated and down-regulated gene sets were carried out with the *Candida* Genome Database Gene Ontology Term Finder (http://www.candidagenome.org/cgi-bin/GO/goTermFinder) using function, process, and component gene ontology (GO) terms. Only hits with a *p* value of <0.05 were considered during the evaluation process (Table S2).

Besides defined GO terms, groups of functionally related genes were also generated by extracting data from the *Candida* Genome Database (http://www.candidagenome.org) unless otherwise indicated (Table S3). The enrichment of *C. albicans* genes from these gene groups in the up-regulated and down-regulated gene sets was tested with Fisher’s exact test (*p* < 0.05). The following gene groups were created:

i. *Virulence-related genes* – Genes involved in the genetic control of *C. albicans* virulence were collected according to Mayer et al. (2013), Höfs et al. (2016) and Araújo et al. (2017) (31–33).
ii. *Antioxidant enzyme-related genes* – This gene group includes genes encoding functionally verified and/or putative antioxidant enzymes according to catalase (GOID:4096), SOD (GOID:4784), glutaredoxin (GOID:6749), thioredoxin (GOIDs:8379 and 51920) and peroxidase (GOID:4601) GO terms.
iii. *Metabolic pathway-related genes* – This group contains all genes related to the carbohydrate, ergosterol and fatty acid biochemical pathways according to the pathway databases (http://pathway.candidagenome.org/).
iv. *Metal metabolism-related genes* – Genes involved in iron, zinc, copper and manganese acquisition by *C. albicans* were grouped according to Fourie et al. (2018) and Gerwien et al. (2018) (34–35).

### Data availability

The data shown and discussed in this paper have been deposited in the NCBI Gene Expression Omnibus (GEO) (36) (https://www.ncbi.nlm.nih.gov/geo/) with the following accession no.: GSE199383

## Results

### Effects of surfactin on growth, morphology, and extracellular phospholipase, proteinase and esterase production by *Candida albicans*

The growth of *C. albicans* was examined following preculturing for 4 hours and subsequently treating with 128 mg/l surfactin in YPD medium. Growth was significantly inhibited within 1 hour after the addition of surfactin as assessed both by observed absorbance values (0.96±0.04 and 0.75±0.02 for untreated control and surfactin-exposed cells, respectively, at OD_640_) (*p*<0.001) (Figure 1A) and DCM changes (0.95±0.1 g/l and 0.6±0.15 g/l for untreated control and surfactin-treated cells, respectively) (*p*<0.01) (Figure 1B). The observed growth inhibition was further confirmed by changes in measured DCM after 6 hours (1.5±0.2 g/l and 0.7±0.1 g/l) and 8 hours of incubation (3.2±0.3 g/l and 1.0±0.2 g/l) for untreated control and surfactin-exposed cells, respectively (*p*<0.01 to 0.001) (Figure 1B).

**Figure 1.**
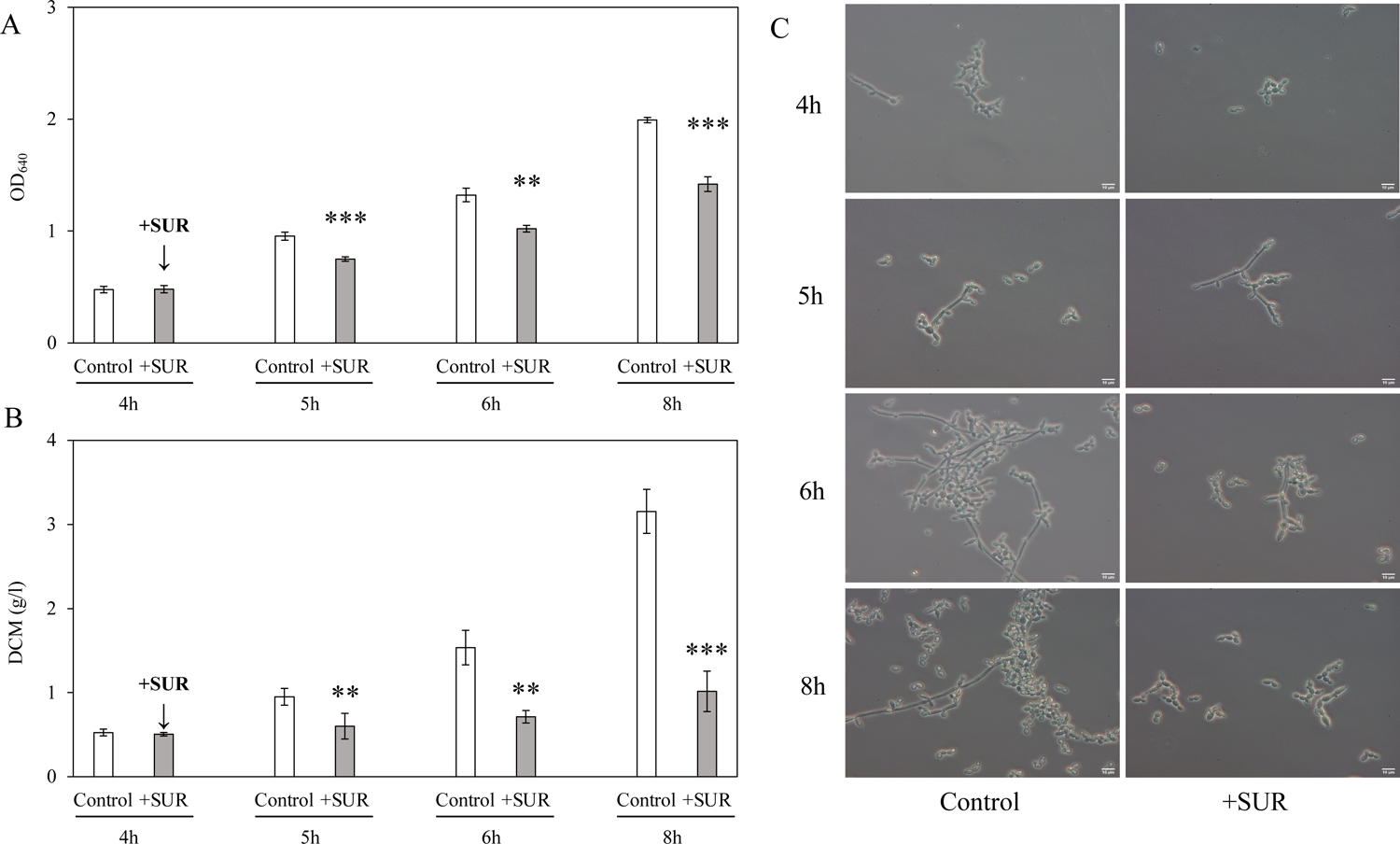
Effect of surfactin on the growth and morphology transition of *Candida albicans*. Changes in the growth of *C. albicans* were examined by measurement of the absorbance (OD_640_) (A) and dry cell mass (DCM) (B). Typical morphological forms can be observed in untreated and surfactin-exposed (SUR) *C. albicans* cultures at 4, 5, 6 and 8 hours (C). Following a 4-hour incubation, surfactin (SUR) was added at a final concentration of 125 mg/l to the YPD cultures. Data represent mean values with standard deviations (SD) calculated from three independent experiments. The asterisks indicate a statistically significant difference between control and surfactin-treated cultures calculated using the paired Student’s *t* test (** *p* < 0.01, *** *p* < 0.001).

The type and ratio of filamentous morphological forms (hypha or pseudohypha) differed significantly compared to the yeast form in treated and untreated cultures. The ratio of hyphae and pseudohyphae was statistically comparable at 5 and 6 hours both for surfactin-treated and control cultures. However, the ratio of hyphae was significantly lower in surfactin-exposed culture compared with untreated cells from 8 hours of incubation (10.33%±2.08% vs. 14.33%±1.53%) (*p*<0.05). Furthermore, the ratio of pseudohyphae was 29.66%±3.06% and 22.67%±3.05% at 8 hours for treated and untreated cells, respectively (*p*<0.05) (Figure 1C).

Testing virulence-related enzymes revealed that surfactin treatment did not significantly influence the extracellular phospholipase (Pz values were 0.6±0.05 and 0.57±0.04), the aspartate proteinase (0.78±0.09 and 0.77±0.09) or the esterase activity (0.63±0.04 and 0.67±0.06) of *C. albicans* when compared with the untreated control cells.

### Surfactin exposure significantly influences glucose consumption, ethanol production, and the intracellular metal contents of *Candida albicans* cells

The concentration of glucose consumption and ethanol fermentation of *C. albicans* was determined for 2 and 4 hours subsequently after surfactin exposure. Surfactin treatment caused a significant increase in glucose utilization at 4 hours post-exposure (4.1±0.3 g/g vs. 2.95±0.4 g/g DCM) (*p*<0.001) (Figure 2A). Enhanced ethanol fermentation was observed both at 2 hours (7.8±2.2 g/g DCM vs. 0.3±0.1 g/g DCM) (*p*<0.01) and 4 hours post-exposure (5.1±0.7 g/g DCM vs. 0.64±0.08 g/g DCM) (*p*<0.001) (Figure 2B).

**Figure 2.**
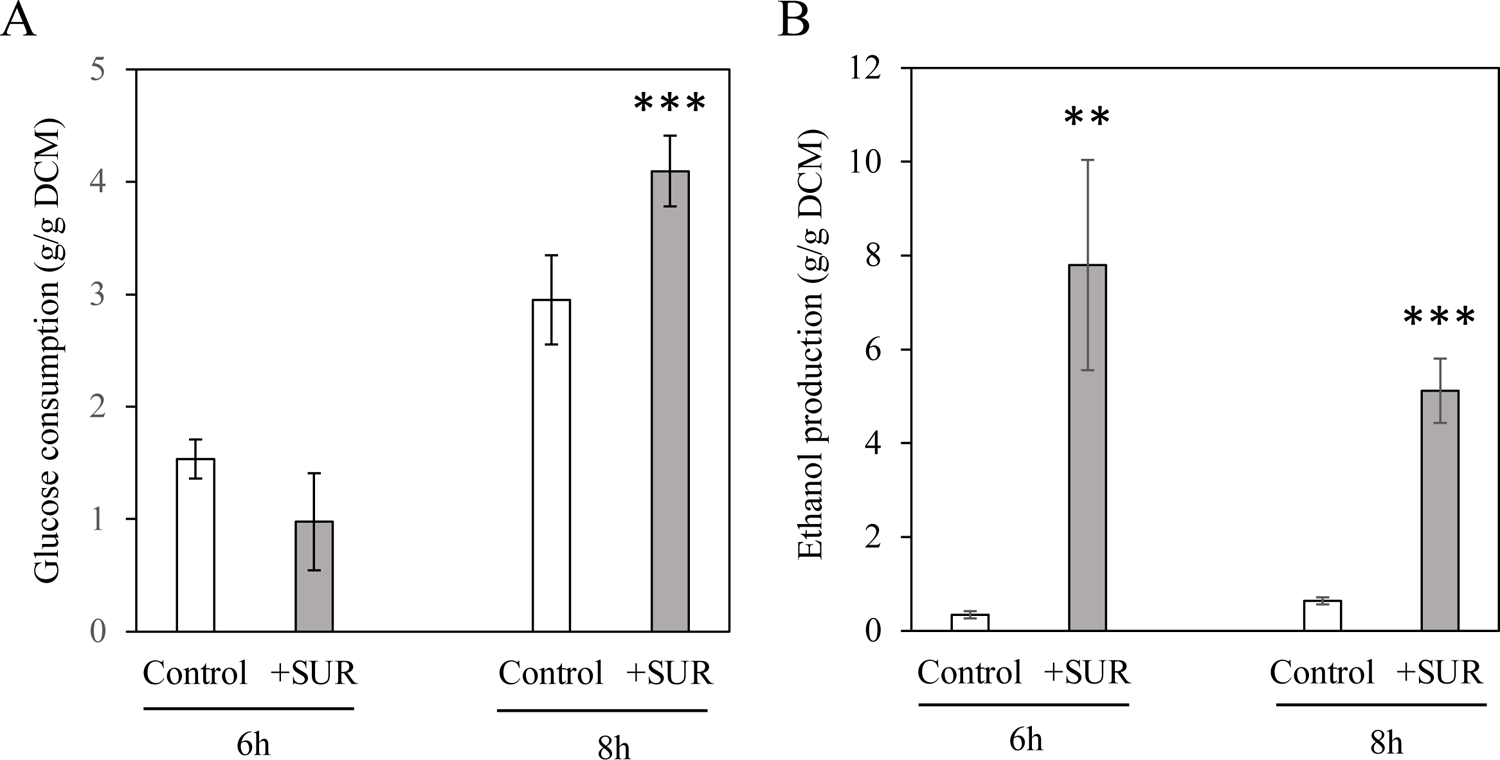
The physiological effect of surfactin exposure in *Candida albicans*. The physiological effects of surfactin exposure on glucose utilization (A) and ethanol production (B). Cell cultures were exposed to 128 mg/l surfactin (SUR) for 4 hours. Samples were taken after 2 and 4 hours of surfactin exposure. Data represent mean values ± standard deviation calculated from three independent experiments. *, ** and *** indicate significant differences at *p* values of <0.05, 0.01 and 0.001, respectively, calculated using the paired Student’s *t* test, comparing untreated control and surfactin-treated cultures.

Following 4 hours of surfactin treatment, the iron, manganese and zinc content of yeast cells decreased (70.9±8.4 mg/kg vs. 127.1±18.4 mg/kg, 5.3±1.3 mg/kg vs. 8.1±0.5 mg/kg, and 169.6±8.9 mg/kg vs. 214.4±4.5 mg/kg, for iron, manganese, and zinc in treated versus non-treated cells, respectively) (*p*<0.05 to 0.01), while the copper content of *C. albicans* cells did not differ significantly from the untreated control cultures (3.1±0.95 mg/kg vs. 2.0±0.25 mg/kg) (*p*>0.05) (Table 1).

**Table 1.** Surfactin-induced iron, manganese, zinc, and copper content in *Candida albicans*

| Metal contents/<br>Treatment | Mean value± SD <sup>a</sup> |  |  |  |
| --- | --- | --- | --- | --- |
|  | Fe<br>(mg/kg) | Mn<br>(mg/kg) | Zn<br>(mg/kg) | Cu<br>(mg/kg) |
| Untreated cultures | 127.1±18.4 | 8.1±0.5 | 214.4±4.5 | 2.0±0.25 |
| Surfactin-treated cultures | 70.9±8.4 * | 5.3±1.3 * | 169.6±8.9 ** | 3.1±0.95 |

<sup>a</sup> Mean values ± standard deviations (SD) calculated from three independent experiments are presented.
\*and \*\* indicate significant differences at *p* values of <0.05 and 0.01, respectively, calculated by paired Student's *t* test, comparing untreated control and surfactin-treated cultures.

### Surfactin exposure increased the intracellular glutathione concentration of *Candida albicans* cells

Surfactin treatment did not enhance oxidative stress in *C. albicans* cells. DCF production as a marker of redox imbalance in the surfactin-treated cultures did not differ significantly from that observed in untreated control cultures (12.9±2.7 nmol DCF/OD_640_ and 11.5±1.8 nmol DCF/OD_640_, respectively) (*p*>0.05). Nevertheless, the GSH concentration was significantly higher in the surfactin-challenged cells (37.2±10.4 nmol/mg vs. 2.05±0.8 nmol/mg) (*p*<0.01) than in untreated control cells, while surfactin treatment did not result in a significantly higher GSSG level (0.05±0.01 nmol/mg vs. 0.03±0.007 nmol/mg) (*p*>0.05).

### Surfactin exposure inhibits the adhesion of *Candida albicans* to inert surface and to Caco-2 cells

The metabolic activity of inert surface-adhered *C. albicans* cells is depicted in Figure S1. In the first 12 hours, the relative metabolic activity of surfactin-exposed cells was significantly lower than that of the untreated control (*p*<0.01-0.001). Noticeably, the lowest metabolic activity was observed at 6 hours (48.07%±4.25%).

Adhesion of fungal cells to Caco-2 cells was significantly lower already 1 hour post surfactin exposure (2.2%±0.8% adherent cells) compared with untreated control cells (5.8%±1.1% adherent cells) (*p*<0.01).

### Surfactin enhances the *in vitro* activity of fluconazole against *Candida albicans*

The nature of the interaction between surfactin and fluconazole was evaluated using FICI calculation and the BLISS independence model. Based on three independent experiments, the median MIC values for fluconazole and surfactin were 0.125 mg/l (0.125–0.25 mg/l) and >256 mg/l (256 to >256 mg/l) when added separately, while 0.004 mg/l and 8 mg/l (8–32 mg/l) in combination, respectively. A synergistic interaction between fluconazole and surfactin was observed for all three independent experiments, with FICI ranging from 0.135 to 0.37. The FICI values were clearly confirmed by the Bliss independence model, where the mean synergy score was 22.7 (Figure S2).

### Transcriptional profiling and RNA-Seq data validation

Principal component analysis (PCA) and hierarchical clustering were used to represent visually the transcriptomic differences between samples treated with surfactin and the untreated controls. Analyses of the RNA sequencing data clearly indicated that surfactin has a remarkable effect on *C. albicans* gene expression, leading to significant alterations in the transcriptome (Figure S3).

Comparison of the surfactin-treated *C. albicans* global gene expression profile with that of unexposed cells revealed 1389 differentially expressed genes. Among those, 773 were up-regulated and 617 were down-regulated in the surfactin-exposed samples compared to the untreated controls (Figure 3A–C, Figure 4, Tables S2 and S3).

**Figure 3.**
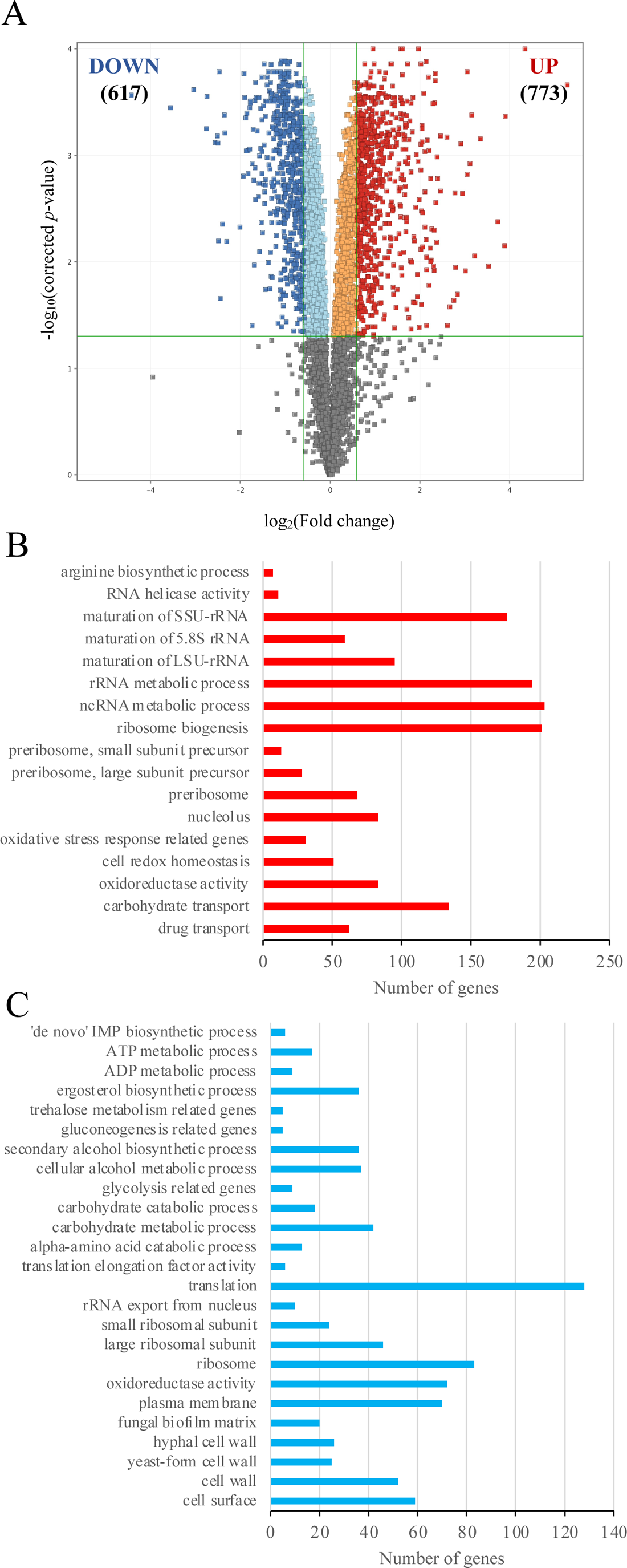
Overview of transcriptional changes induced by surfactin in *C. albicans*. Up-regulated (red) and down-regulated (blue) genes were defined as differentially expressed genes (corrected *p* value of <0.05), with more than a 1.5-fold increase or decrease in their transcription (surfactin treated versus untreated) (A). Summary of gene enrichment analyses and the number of genes affected by *C. albicans* exposure to surfactin (B–C). The enrichment of up-regulated (B) and down-regulated (C) gene groups was identified using the Candida Genome Database Gene Ontology Term Finder (http://www.candidagenome.org/cgi-bin/GO/goTermFinder) or was tested by Fisher’s exact test. The data sets for the gene groups are available in Tables S2 and Table S3 in the supplemental material.

**Figure 4.**
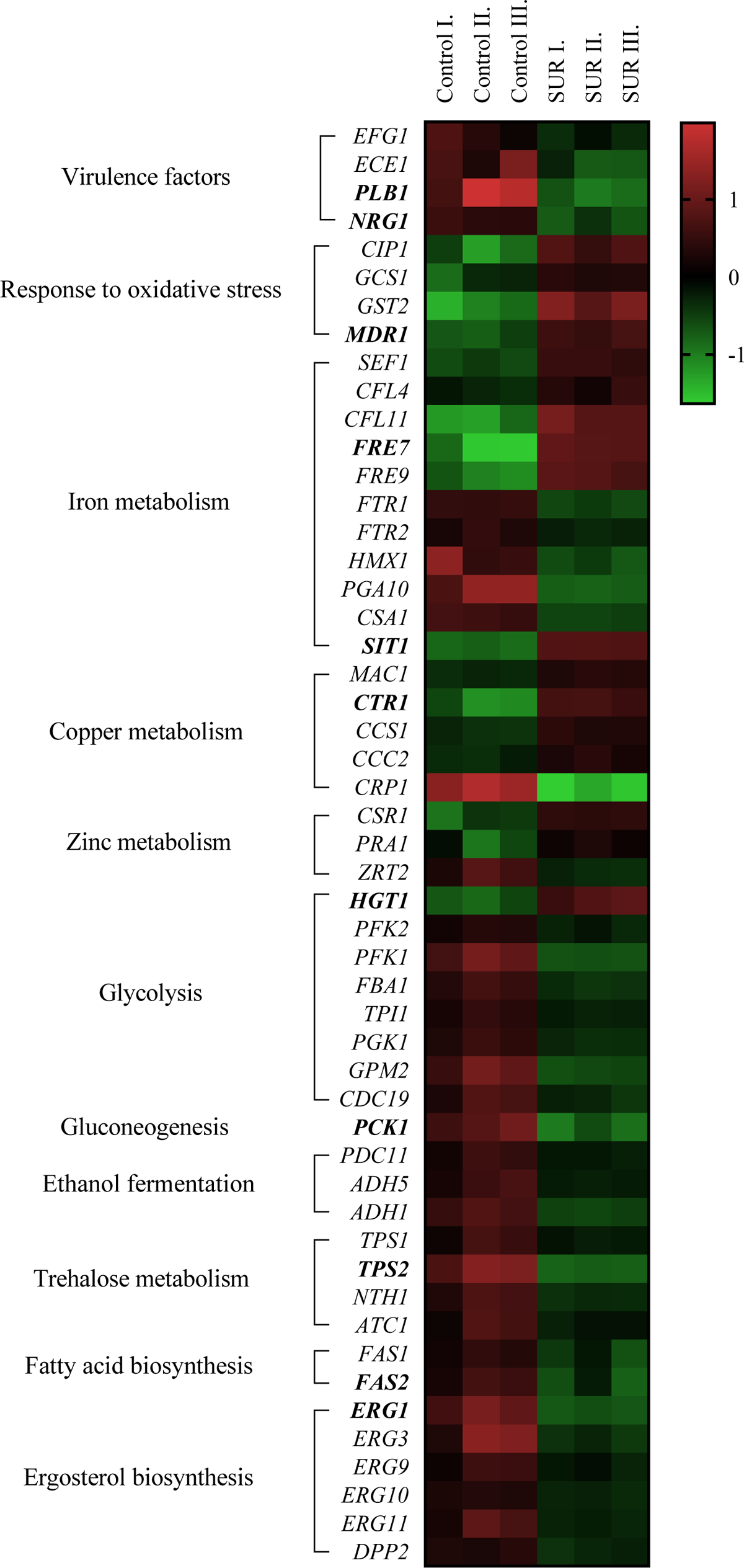
The effects of surfactin on the expression of selected genes of *C. albicans*. The heat map demonstrates the expression profiles of representative genes according to the colour scale that indicates gene expression changes. Bold names indicate the genes that were selected for analysis by RT-qPCR. The data sets for the gene groups are available in Tables S2, Table S3 and Table S4 in the supplemental material.

### Evaluation of surfactin-responsive genes

To obtain an overall comprehensive insight into the molecular effect of surfactin, gene ontology terms were assigned to all of the genes in the *C. albicans* reference genome; after which, we compared the terms for both the down-regulated and up-regulated genes with a background of all terms.

#### Virulence-related genes

In agreement with reduced surface adhesion (see above), biofilm-related genes were significantly enriched within the surfactin-responsive down-regulated gene group according to Fisher’s exact test (Table S3). Nine genes are involved in biofilm adhesion (*EAP1* and *PGA1*), biofilm maturation (*EFG1*, *ROB1, CPH2*, *IFD6*, *GCA1,* and *ADH5*), and biofilm dispersion (*NRG1*) (Figure 3A–C, Figure 4, Table S3). Down-regulation of *PLB1* and *NRG1* expression following surfactin treatment was supported by RT-qPCR data (Figure 4 and Table S4).

#### Oxidative stress-related genes

Surfactin exposure increased the expression of glutathione metabolism-related genes (*GTT11, GTT12, GTT13*, *GCS1*, *GST2*, *C5_01560C*, and *TTR*1) and flavodoxin-like protein (FLPs) genes (*PST1*, *PST2*, and *C2_07070W*) as well as the *CIP1* gene, which encodes an environmental stress-induced protein (Figure 3A–C, Figure 4, Table S3).

#### Metabolic pathway-related genes

Surfactin treatment led to decreased expression of several genes involved in glycolysis (*PFK2*, *PFK1*, *FBA1*, *TPI1*, *PGK1*, *GPM2*, *CDC19*, *HXK2*, and *GAL4*), ethanol fermentation (*PDC11, ADH5,* and *ADH1*), trehalose metabolism (*TPS1, TPS2, TPS3, NTH1,* and *ATC1*), glycogen biosynthesis (*C1_01360C, GSY1, GLC3,* and *PGM2*), gluconeogenesis (*MAE1, PCK1, GPM2, PGK1*, and *FBA1)*, ergosterol biosynthesis (*ERG1, ERG3, ERG9, ERG10,* and *ERG11*) and fatty acid biosynthesis (*ACC1, FAS1,* and *FAS2*) (Figure 3A–C, Figure 4, Table S3).

In addition, surfactin exposure caused a significant increase in the expression of preribosome biogenesis-related genes, including small subunit genes (*BUD21*, *BUD23*, *RPS27A*, *SAS10*, *DBP8* etc.) and large subunit genes (*CSI2*, *DBP3*, *SDA1*, *SPO22*, *YTM1* etc.)

Genes involved in iron homeostasis including ferric reductases (*CFL4, CFL11, CR_07300W, C7_00430W, FRE7,* and *FRE9)*, multicopper oxidases (*FET99* and *CCC2*), siderophore transport *(SIT1*), and *SEF1* transcription factor as well as copper homeostasis (*MAC1* encoding transcription factor*, CCS1, CTR1,* and *CCC2* coding for chaperone and copper transporters) and zinc metabolism (*CSR1* and *PRA1* encoding transcription factor and cell surface protein) were enriched in the up-regulated gene set (Figure 3A–C, Figure 4, Table S3).

In addition, surfactin exposure decreased the expression of ferric permease genes (*FTR1* and *FTR2*), hemoglobin utilization-related genes (*HMX1, PGA10*, and *CSA1*) and the *AFT2* gene encoding a transcription factor as well as vacuolar iron (*SMF3*), copper (*CRP1*) and zinc (*ZRT1*) transporters (Figure 3A–C, Figure 4, Table S3).

The up-regulation of *FRE7*, *SIT1*, and *CTR1* and the down-regulation of *TPS2, PCK1, ERG1,* and *FAS2* were additionally confirmed by RT-qPCR (Figure 4, Table S3).

#### Transmembrane transport-related genes

Surfactin treatment led to increased expression of several genes involved in transmembrane transport, including 62 antifungal drug transporter (e.g. *MDR1* and *FLU1*) and 134 carbohydrate transporter (*HGT1, HGT10, HGT14, HGT18, HXT5, NAG3, SFC1, GAL102* etc.) genes (Figure 3A–C, Figure 4, Table S3).

In addition, 10 genes related to rRNA export from the nucleus (*RPS3, RPS5, RPS10, RPS15, RPS18, RPS19A, RPS21, RPS26A, RPS28B,* and *YST1*) were significantly enriched in the down-regulated gene set (Figure 3A–C, Figure 4, Table S3).

From this above list, RT-qPCR measurements demonstrated that surfactin exposure caused a significant increase in the expression of *MDR1* (ABC transporter) and *HGT1* (glucose transmembrane transporter) in the treated cells (Figure 4 and Table S4).

## Discussion

Commensal co-existence of *C. albicans* with its human host has been a well-known phenomenon for decades. However, several studies have demonstrated that gastrointestinal *C. albicans* colonization is one of the most important predisposing factors for life-threatening invasive *Candida* infections (1–3). Biosurfactant production by probiotic bacterial species or its use as an adjuvant in complementary therapy may prevent the overgrowth of *C. albicans* cells, which could significantly reduce the need for antifungal agents and the risk of resistance development (37). Based on recent *in vitro* findings, one of the most effective biosurfactants is surfactin, which was first described from *B. subtilis* ATCC 21332 and has potent antimicrobial, antitumoral and anticoagulant activities (38). Studies of surfactin focusing on anti-*Candida* activity are scarce; nevertheless, Biniarz et al. (2015) reported that the cell surface hydrophobicity and adhesion of *C. albicans* decreased by 20–60% and 80–90%, respectively following surfactin exposure (39). Similar results were observed with regard to the inhibition of adherence and biofilm development by Nelson et al. (18). However, the detailed physiological and transcriptional background of the preliminary antifungal activity caused by surfactin remained to be elucidated.

Our work reveals several roles for *B. subtilis* surfactin as a potent modulator of *C. albicans* metabolism, morphogenesis and susceptibility to traditional antifungal drugs. The detected growth inhibition can be attributed to the general down-regulation of numerous metabolism-related genes. Although glucose consumption was significantly higher in surfactin-exposed cells at 4 hours, prior to this time this value was lower compared with the control, which coincided with the measured reduction of metabolic activity in the case of surfactin-treated adhered fungal cells. Interestingly, ribosome biogenesis-related genes were significantly up-regulated by surfactin treatment. Ribosome biogenesis is closely linked to such cellular activities as growth and division (40). After transcription, both the small and large pre-ribosomal subunits must be exported to the cytoplasm to allow ribosome maturation. Our GO term enrichment analysis revealed that surfactin treatment up-regulates the transcription of genes connected to RNA metabolism (220 genes) including the pre-ribosome-related genes; furthermore, it down-regulates the genes of rRNA transport (10 genes), cytosolic ribosomal proteins (69 genes) and translation (128 genes). These transcriptomic results suggest that the reduced growth of *C. albicans* cells can be also attributed to the suspension of protein synthesis under surfactin treatment (40) in addition to altering expression of genes connected to metabolism.

Several ergosterol (*ERG1*, *ERG3*, *ERG9*, *ERG10*, *ERG11*) and fatty acid metabolism-related genes (*ACC1*, *FAS1*, *FAS2*) were down-regulated, suggesting possible changes in the structure and fluidity of the cell membrane. The observed synergistic interaction between surfactin and fluconazole supports these presumed membrane changes (41). Surprisingly, a remarkable increase in ethanol content was detected in surfactin-exposed cultures both at 2 and 4 hours compared to the untreated control. Similarly, *Pseudomonas aeruginosa*-derived phenazines increase the production of fermentation products such as ethanol by 3- to 5-fold in *C. albicans* (42). Presumably, surfactin possibly shapes the chemical ecology of mixed-species microbial communities. *C. albicans* biofilm development may also be influenced by this elevated ethanol content, which has been shown to inhibit biofilm formation (43). Moreover, transcription of both *EFG1* and *ECE1* were down-regulated, explaining the inhibition of hyphae formation and biofilm development (44). A surfactin-diminished biofilm production may potentially explain its probable interfering effect with fungal quorum sensing. A previous studies highlighted that surfactin might act as a signal molecule in a paracrine signaling cascade of *B. subtilis* (45). Furthermore, surfactin might also be involved in interaction of *B. subtilis* with the black mold fungus, *Aspergillus niger*, resulting in altered metabolism both in the bacterium and the fungus (46). Our data suggest that surfactin may significantly influence the farnesol-mediated quorum sensing pathways in *C. albicans* because farnesol synthesis was disturbed by the surfactin-induced down-regulation of the lipid pyrophosphatase enzyme encoding *DPP2* (47). Tyrosol production may also be disrupted based on the down-regulated level of *ADH1* and *ADH5*, both of which are involved in tyrosine-tyrosol conversion in *C. albicans*, further explaining the observed inhibition of biofilm formation (48). Regarding the inter-species interaction on biofilm formation, Liu et al. (2019) reported the extensive inhibition of quorum sensing-related pathways by surfactin, including biofilm formation by *Staphylococcus aureus* (49). Based on their findings, surfactin has a significant effect on polysaccharide production and decreases the level of alkali-soluble polysaccharide in biofilms (49).

The here obtained transcriptomic data highlighted that surfactin treatment significantly influenced the transcription of iron metabolism, copper transport and zinc transport-related genes as well as the intracellular iron, zinc and manganese contents of *C. albicans*. Notably, similar consequences were observed in the case of *C. auris* following farnesol exposure (23). The decreased iron content may be part of the general defense against surfactin to minimize the damage caused by ferrous ions (50). In previous studies, the elevated level of free intracellular iron facilitated the production of reactive oxygen species and triggered iron-dependent cell death in *Saccharomyces cerevisiae* (51). The surfactin-induced reduction in intracellular zinc content was associated with the down-regulation of the *ZRT2* gene, which encodes the major zinc importer in *C. albicans*. Zinc is a crucial essential element in oxidative stress defense as a structural component of superoxide dismutase; furthermore, *ZRT2* deletion or down-regulation impaired virulence by decreasing the colonization rate of *C. albicans* and reducing kidney fungal burden in a mouse infection model (52).

Copper homeostasis is also an important determinant for virulence in *C. albicans*. In our work, transcription of *CTR1* was up-regulated simultaneously with the *FRE7* gene. Khemiri et al. (2020) reported that under copper deprivation, *C. albicans*-activated genes required for copper uptake included the transporter Ctr1 and the ferric reductases Fre7 (52). At the same time, transcription of *CRP1* was down-regulated, which encodes a P-type ATPase that functions as a copper extrusion pump to facilitate survival in high copper environments (52). We hypothesize that surfactin binds to Cu^2+^ ions, mimicking copper deprivation. A previous study showed that Cu^2+^ ions preferentially bind to amide nitrogens of lipopeptide biosurfactants such as *B. subtilis*-derived surfactin (53).

Based on our data, surfactin did not generate elevated levels of reactive oxygen species. Nevertheless, GSH was significantly increased upon surfactin treatment. Studies in *C. albicans* suggest involvement of GSH in the yeast to mycelium morphological switching (54). The elevated GSH level is associated with inhibition of hyphae formation. Furthermore, the overproduction of GSH may induce reductive stress response explaining partly the observed inhibition (54). Presumably, surfactin can become conjugated to GSH during attempts to inactivate it and remove it from fungal cells, thus minimizing the damage caused by this compound. Importantly, the transcription level of the gene encoding the Mdr1-P-glycoprotein was significantly up-regulated, further suggesting its role in surfactin detoxification (55).

In summary, this is the first study examining the changes in gene transcription in *C. albicans* following surfactin exposure. Surfactin decreased the adherence of fungal cells, modulated the morphology toward yeast and pseudohyphae forms, and inhibited biofilm formation. Moreover, surfactin significantly repressed the expression of ergosterol synthesis-related genes and major metabolic pathways. It also reduced iron, manganese and zinc metabolism, but significantly enhanced ethanol production. Focusing on its potential applicability as an adjuvant or probiotic-derived compound, the most important advantages of surfactin are its low risk of toxicity, high biodegradability properties and long-term physico-chemical stability. Nevertheless, two major limitations should be highlighted. Biotechnological processes including the synthesis of surfactin are currently relatively expensive and optimization of the purification process may be problematic. Furthermore, the extent of the available data focusing on *in vivo* experiments is limited. Nonetheless, there is huge potential for surfactin and surfactin-producing microbes, especially against gastrointestinal candidiasis, and it would be a mistake to leave this potential untapped.

## Ethical approval

Not required

---

**Supplementary Figure S1 Changes in metabolic activity over time in *Candida albicans* adhered to an inert surface in the presence of 128 mg/l surfactin.**

The untreated control cells showed 100% metabolic activity at each time point. Data represent mean values of relative metabolic activity ± standard deviation calculated from three independent experiments. Statistical analysis was performed using the paired Student’s *t* test.

**Supplementary Figure S2 The nature of interaction using the BLISS independence model**

2D (A) and 3D (B) synergy maps demonstrate the combination landscape of fluconazole with surfactin. Synergy maps were generated using the open access “SynergyFinder” program.

**Supplementary Figure S3 Cluster (A) and principal component (B) analysis of the transcriptome data.**

Symbols represent untreated control (Cont) and 128 mg/l surfactin-exposed (SUR) cultures. The distribution of transcriptome data obtained in three independent series of experiments (I, II, and III). Analyses were performed with the StrandNGS software using default settings.

**Supplementary Table S1 Oligonucleotide primers used for RT-qPCR analysis.**

**Supplementary Table S2 Results of the gene set enrichment analysis.**

Significant shared GO terms (*p* < 0.05) were determined with the Candida Genome Database Gene Ontology Term Finder (http://www.candidagenome.org/cgi-bin/GO/goTermFinder). Up- and down-regulated genes were defined as differentially expressed genes where log_2_(FC) > 0.585 or log_2_(FC) < −0.585. The FC ratios were calculated from the normalized gene expression values. Biological processes, molecular function and cellular component categories are provided.

**Supplementary Table S3 Transcription data of selected gene groups.**

Part 1: Genes involved in genetic control of *Candida albicans* virulence.

Part 2: Genes involved in antioxidative defense.

Part 3: Genes involved in selected metabolic pathways.

Part 4: Genes or ergosterol and fatty acid metabolism.

Part 5: Genes involved in iron, zinc, copper and manganese metabolism.

Part 6: Genes involved in autophagy.

Part 7: Genes involved in apoptosis.

The systematic names, gene names and the features (putative molecular function or biological process) of the genes are given according to the *Candida* Genome Database (http://www.candidagenome.org).

Up- and down-regulated genes are marked with red and blue colour. Up- and down-regulated genes were defined as differentially expressed genes with >1.5-fold change (FC, up-regulated genes) or less than −1.5-FC (down-regulated genes) values. The FC ratios were calculated from the normalized gene transcription values.

Results of gene enrichment analysis (Fisher’s exact test) are also enclosed. “The response of oxidative stress” genes (GOID: 0006979) collected from Gene Ontology Term Finder (http://www.candidagenome.org/cgi-bin/GO/goTermFinder).

**Supplementary Table S4 Results of RT-qPCR experiments**

RNA-Seq data are presented as FC values, where FC is “fold change”. Relative transcription levels were quantified as ΔΔCP = ΔCP_control_ − ΔCP_treated_, where ΔCP_treated_ = CP_tested gene_ − CP_reference gene_, measured from surfactin treated cultures, and ΔCP_control_ = CP_tested gene_ − CP_reference gene_, measured from control cultures. CP values represent the RT-qPCR cycle numbers of crossing points. The *HPT1* gene was used as a reference gene. ΔΔCP values significantly (*p* < 0.05 by Student’s *t* test; n = 3) higher or lower than zero (up- or down-regulated genes) are marked in red and blue, respectively. Pearson’s correlation coefficient between the RT-qPCR and RNA-Seq values was 0.95.

